# Deep Learning Reveals Cross-Modal Neural Representations of Auditory and Visual Mental Imagery in MEG

**DOI:** 10.64898/2026.02.02.703195

**Authors:** Alina Schüller, Constantin Jehn, Marietta Stegmaier, Jasmin Riegel, Tobias Reichenbach

**Affiliations:** Department Artificial Intelligence in Biomedical Engineering (AIBE), Friedrich-Alexander-Universität Erlangen-Nürnberg (FAU), Nürnberger Straße 74, 91052 Erlangen, Germany

**Keywords:** CNN, imagined speech, MEG, memory, neural decoding, visual imagery

## Abstract

Mental imagery provides a unique window into the brain’s ability to internally simulate sensory experiences, offering valuable insights for both cognitive neuroscience and brain–computer interface (BCI) research. This study examined the neural representations of imagined auditory and visual stimuli using magnetoencephalography (MEG) and assessed the ability of machine learning models to decode these mental processes. MEG data were recorded from 18 right-handed participants during auditory and visual imagery tasks and source-reconstructed within modality-specific cortical regions of interest. We compared a convolutional neural network (CNN) and a linear logistic regression model within a subject-specific classification framework. Both approaches achieved above-chance decoding accuracies, with the CNN outperforming the linear model in the auditory task, whereas the linear model showed slightly higher accuracy for visual imagery. Notably, the CNN achieved significant decoding performance even when trained on non–task-relevant cortical regions, indicating that imagined stimuli are represented in distributed and partially overlapping neural networks across modalities. This cross-modal decoding capability highlights the potential of deep learning models to capture complex, multimodal neural patterns and suggests that future brain–computer interfaces could benefit from integrating auditory and visual information. A secondary, behavioral analysis revealed correlation of memory capacity and individual learning preferences with decoding performances, suggesting that individual cognitive differences may further shape the quality of neural representations. Together, these findings advance our understanding of cross-modal mental imagery and point toward more flexible and personalized approaches in BCI design.

**New and Noteworthy:** By comparing linear and deep classifiers, this work shows that convolutional networks capture rich, cross-modal neural representations of auditory and visual mental imagery in MEG. Significant decoding from non-task-relevant regions indicates distributed cortical engagement, highlighting deep learning’s potential for robust, modality-independent brain–computer interfaces.

## Introduction

Neurological disorders such as amyotrophic lateral sclerosis (ALS) or severe brain injury can lead to locked-in syndrome, a condition in which individuals retain full cognitive function but lose voluntary muscle control. For these patients, brain–computer interfaces (BCIs) offer a means of communication and interaction independent of motor output [Dash et al., 2020, van den Berg et al., 2021]. BCIs translate neural activity into control signals for external devices such as communication aids or prosthetic systems, thereby improving autonomy and quality of life.

Conventional BCIs often rely on motor imagery or stimulus-driven paradigms that require sustained attention and extensive user training [Abiri et al., 2019, Rashid et al., 2020]. Additionally, many BCIs require users to spell words letter by letter or imagine movements for every command, leading to high cognitive effort. Most systems also suffer from a low information transfer rate (ITR), making them inconvenient for regular use [Rashid et al., 2020]. As a more natural and intuitive alternative, decoding inner or imagined speech, the silent internal rehearsal of words or phrases, has gained increasing interest [Panachakel and Ramakrishnan, 2021, Bocquelet et al., 2016]. Neural activity during inner speech engages brain regions involved in overt speech perception and production, particularly in temporal and frontal cortices [Amit et al., 2017, Bocquelet et al., 2016], making it a promising signal source for a more seamless and user-friendly communication system.

In parallel, visual imagery, the internal generation of mental images, has also been explored as a complementary modality for BCI control [Kosmyna et al., 2018]. The human brain allocates extensive cortical resources to vision and can extract complex scene information rapidly (e.g., within 150 ms) [Thorpe et al., 1996], making visual imagery a fast and robust channel for representing perceptual content. This efficiency and capacity for detailed visual representation motivate the use of visual imagery as a promising modality for BCI control. Visual imagination therefore offers a stable and intuitive channel for decoding mental content. Importantly, recent neuroimaging studies have shown that auditory and visual imagery are not fully independent processes: visual areas can become active during inner speech, and auditory or linguistic regions may be recruited when processing visual scenes [Amit et al., 2017, Nishimura et al., 2015]. This functional overlap suggests that mental imagery involves distributed and cross-modal neural representations rather than strictly segregated modality-specific networks. Supporting this idea, Lee et al. (2019) reported comparable imagery intensities between auditory and visual tasks, indicating that both modalities may serve as equally viable channels for neural communication.

Several studies have explored the decoding of neural signals and the application of machine learning techniques in brain-computer interfaces. Marion et al. (2021) explored EEG responses to imagined music and demonstrated that imagined neural signals can be predicted accurately through regression analyses. The model outputs were robust enough to allow for precise identification of the imagined musical piece. Similarly, Kosmyna et al. (2018) also analyzed the decoding of electroencephalographic (EEG) signals during imagery tasks, yet focusing on visual instead of auditory imagination. Given that EEG-based BCIs can not only differentiate between visual observation and visual imagery tasks but also distinguish between different observed stimuli, the visual modality shows a promising potential to expand BCI control strategies [Kosmyna et al., 2018, Amit et al., 2017, Lee et al., 2019].

Decoding such distributed activity patterns requires recording methods with high temporal and spatial precision. While EEG has traditionally dominated BCI research due to its portability and applicability in real-world settings [Kilmarx et al., 2024, Nieto et al., 2022, Marion et al., 2021], its limited spatial resolution constrains the identification of distributed neural sources. Magnetoencephalography (MEG) offers a powerful alternative, providing millisecond temporal accuracy and precise spatial localization [Ahlfors and Mody, 2019]. These properties make MEG particularly suited for capturing the complex, time–frequency dynamics of imagined speech and visual imagery [Nishimura et al., 2015]. Findings by Dash et al. (2020), for instance, demonstrated the feasibility of decoding both imagined and spoken phrases directly from non-invasive MEG signals using machine learning approaches. Although MEG necessitates a complex measurement system and a magnetically-shielded room, making it infeasible for real-world BCI applications, it can hence aid in guiding BCI research by identifying suitable neural response patterns.

In the present study, we recorded MEG data while participants performed auditory and visual imagery tasks. To assess the decoding of imagined content, we trained two complementary classifiers: a linear logistic regression model and a convolutional neural network. The CNN was chosen for its ability to extract multiscale spatial–temporal patterns and its demonstrated success in imagined speech decoding [Dash et al., 2020]. Our primary goal was to evaluate how well each model decodes auditory and visual imagery from MEG data and to determine whether decoding generalizes beyond task-relevant cortical regions, thereby testing for cross-modal representations. Additionally, it is a well-known fact that people have diverse cognitive preferences for learning, utilizing distinct sensory modalities to optimize information acquisition and comprehension [Fleming and Mills, 1992]. Hence, we wondered whether preferred learning styles and memory capacities can be quantified using the decoding accuracies of the neural signals. We explored whether decoding accuracy relates to individual cognitive styles, as assessed by the *VARK* questionnaire [Fleming and Mills, 1992] and a modality-specific memorization test. Together, these analyses aim to advance our understanding of distributed neural processing during mental imagery and to inform future BCI designs that leverage multimodal and individualized decoding strategies.

## Materials and methods

### Experimental design and data analysis

#### Data acquisition

A total of 18 healthy, right-handed individuals (9 male and 9 female; 19-31 years) without hearing impairments took part in the study. Participants with visual impairments performed the task with corrected-to-normal vision. The study was granted ethical permission by the ethics board of the University Hospital Erlangen (Registration 22–361-S). The experiment lasted one hour and consisted of auditory and visual stimuli.

Auditory stimuli comprised three German abstract nouns: *Ehre* (honor), *Absicht* (intention), and *Idee* (idea). They were carefully chosen for their similar length (all consisting of two syllables), distinct initial vowels, and low potential to evoke visual associations. Visual stimuli consisted of three color-distinct images: a yellow star, a red heart, and a green-brown tree. The block structure, illustrated in Figure 1, was inspired by earlier imagery research [Nieto et al., 2022, Dash et al., 2020, Kosmyna et al., 2018], and consisted of auditory as well as visual blocks.

**Figure 1.**
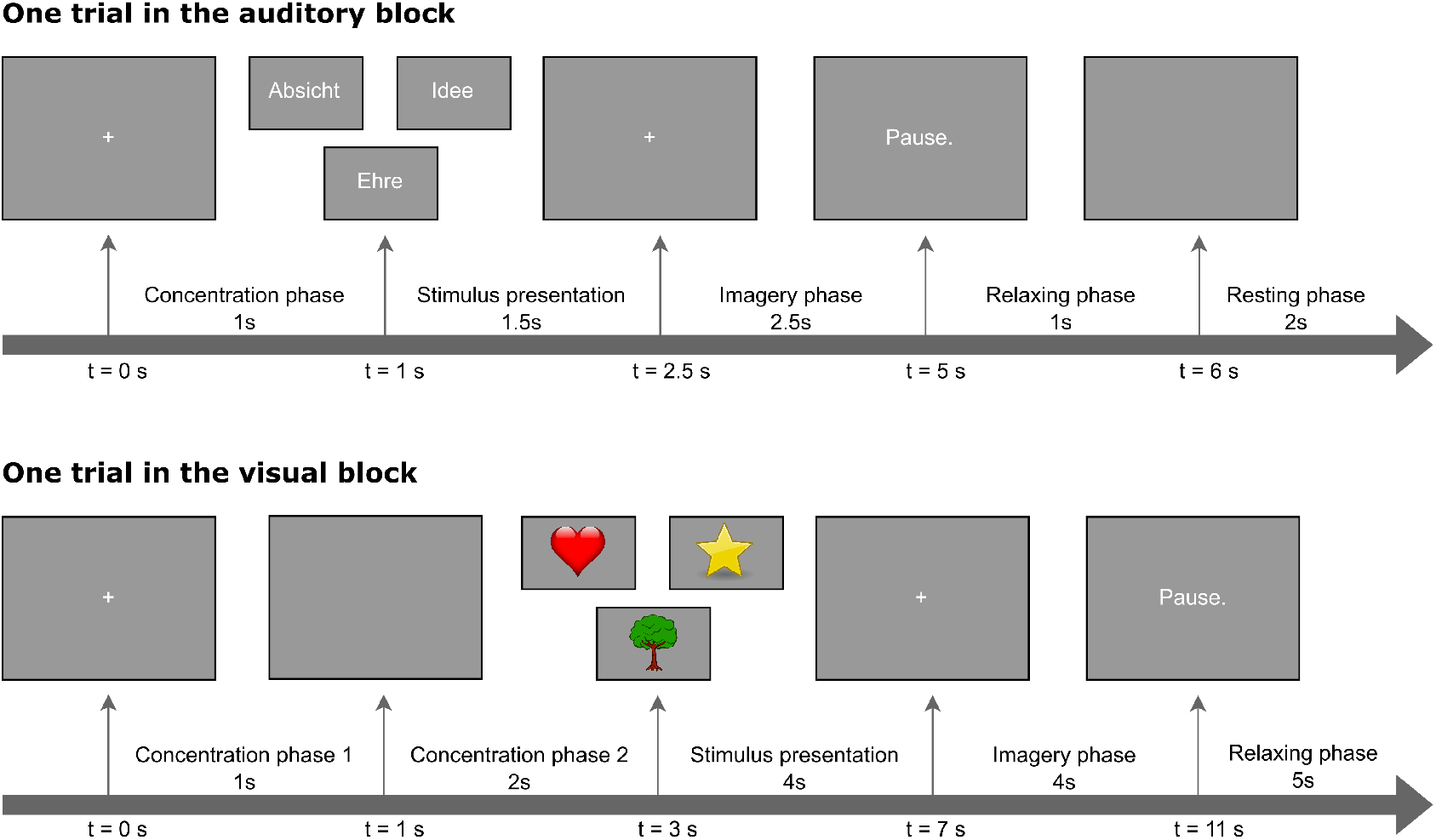
Trial workflow for auditory and visual imagery tasks. Each trial in the auditory block began with a 1 s concentration phase, followed by a 1.5 s word presentation, 2.5 s of auditory imagery, 1 s relaxation, and 2 s rest. Each trial in the visual block began with a 1 s concentration phase, followed by a 4 s image presentation, 4 s of visual imagery, and a 5 s relaxation period. Auditory stimuli consisted of the three German abstract nouns *Ehre* (honor), *Absicht* (intention), and *Idee* (idea). The visual stimuli comprised three color-distinct images (yellow star, red heart, green-brown tree).

In one trial of an auditory block, the participants first entered a 1 s concentration phase to focus on the upcoming sound, followed by a 1.5 s stimulus presentation. They then engaged in 2.5 s of auditory imagery, mentally rehearsing the word they had read. This was followed by 1 s of relaxation and 2 s of rest before the next trial. Each of the three words was presented 15 times per block, yielding 45 trials per block and resulting in 225 auditory trials in total per participant after completing five auditory blocks. The presentation order of the words was randomized in each block.

Each trial in the visual block began with a 1 s concentration phase as well, after which the image was displayed for 4 s. Participants then spent 4 s visualizing the image as vividly as possible while avoiding verbal associations with the corresponding object name (*star, heart, tree*). A 5 s relaxation period followed before the next trial. Each image was shown 6 times per block, resulting in 18 trials per block and 90 visual trials in total per participant after completing the five visual blocks. The presentation order of the images was randomized in each block.

Each session consisted of five auditory and five visual blocks that were presented alternatingly. For half of the participants, the session began with an auditory block followed by a visual block and so on; the remaining participants started with a visual block. Short breaks were provided between blocks.

MEG recordings were obtained with a 4D Neuroimaging system (248 magnetometers) at the University Hospital Erlangen, following the technical specifications conducted within an earlier study by Schüller et al. (2024). Three broken channels were excluded, yielding 245 channels for analysis. Data were recorded at a sampling frequency of 1,017.25 Hz with participants lying supine, eyes open, and an analogue high-pass filter at 1 Hz was applied online. A Polhemus-integrated digitizer recorded five anatomical landmarks. Environmental noise was suppressed using the manufacturer’s calibrated linear weighting algorithm on 23 reference sensors. A stimulation computer interfaced with an external USB audio device providing five analogue outputs; two fed an audio amplifier, and one was additionally routed to an MEG analogue input channel. To mark imagery onset and offset, inaudible 300 Hz, 0.01 s trigger tones were generated and recorded via an analogue audio reference channel, enabling temporal alignment of the imagined intervals with MEG signals. The participants’ heads were recorded with a camera and monitored outside the MEG chamber to control for head and facial movements. Electrooculogram (EOG) activity was recorded to monitor facial and lip movements and no interference signals were observed.

Finally, participants underwent Magnetic Resonance Imaging (MRI) to facilitate individual source reconstruction based on each participant’s anatomical template. T1-weighted MR images were acquired using a Siemens 3T Magnetom Cima.X scanner.

#### Learning preferences and memorization tasks

Apart from MEG acquisition, on a different day, participants completed visual and auditory memorization tasks. In the visual task, a 6×6 grid of white squares was presented on a computer screen. In each trial, seven randomly chosen squares were sequentially highlighted in red, after which participants were asked to reproduce the sequence by clicking them in correct order. In the auditory task, subjects heard a sequence of seven syllables randomly drawn from a set of 35, presented via earphones by an AI-generated female voice. Participants reproduced the sequence by selecting the correct syllables from a displayed grid. Both tasks were performed five times resulting in five trials. Performance was quantified through the auditory memorization score 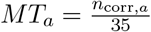 for the auditory memorization task, and the visual memorization score 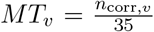 for the visual memorization task. *n*_corr,*a*_ and *n*_corr,*v*_ are hereby the number of correctly recalled syllables and squares, respectively.

All stimulus presentations described above, including the stimulus sequence during the MEG session and the memorization tasks, were programmed in PsychoPy, an open-source Python package for designing and running experiments in behavioral and cognitive neuroscience [Peirce et al., 2019].

In addition to the memorization tasks, participants completed the *VARK* questionnaire, developed by Fleming and Mills (1992), which assesses individual learning preferences through 16 questions, with each offering four response options corresponding to four modalities: visual (*v*), auditory (*a*), read/write (*r*), and kinesthetic (*k*). The resulting four *VARK* scores were calculated for each participant as follows: 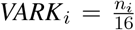, where *n*_*i*_ denotes the number of selected response options and *i* = *a, v, r*, or *k* denotes auditory, visual, read/write or kinesthetic, respectively. Table 1 depicts the scores of the memorization tasks and *VARK* questionnaire, averaged across subjects.

**Table 1:**
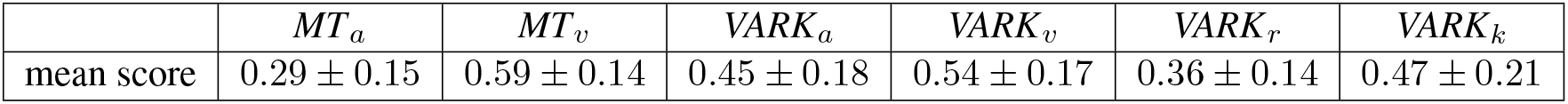
Mean scores (± standard deviation of the mean) of the memorization tasks and *VARK* questionnaire. The subject-averaged auditory and visual memorization task scores *MT*_*a*_ and *MT*_*v*_, respectively, and the four *VARK* modality scores (auditory (*a*), visual (*v*), read/write (*r*), and kinesthetic (*k*)) are presented.

#### Source reconstruction

The acquired MR scans were processed using FreeSurfer [Fischl, 2012], and an *aparc+aseg* volume parcellation was applied to the brain volumes. To localize the neural signals spatially, the raw MEG data were source-reconstructed using an LCMV beamformer onto regions of interest (ROIs) defined from each participant’s individual MRI template. Two separate cortical ROIs, one auditory and one visual, were created to target modality-specific processing. The subject-specific data were finally morphed to the *fsaverage* brain template by FreeSurfer. By that, the auditory ROI comprised 202 source points and, due to anatomical differences of the auditory and visual cortices, the visual ROI contained a smaller set with 107 source points.

#### Preprocessing

MEG data preprocessing was carried out in MNE-Python [Gramfort et al., 2013, Larson et al., 2024]. A 50 Hz notch filter was applied to remove power line noise and the data were resampled to 1,000 Hz. Individual imagery trials were segmented from the continuous MEG signal using the trigger timestamps recorded in the reference channel. The imagination of the three auditory stimuli did not occupy the full 2.5 s trial period, thus, retaining the entire window introduce substantial noise detrimental to classifier performance. Pronouncing the longest words included in the stimuli (*Absicht*), takes around 1 to 1.5 s. Therefore, only the first 1.2 s of the auditory imagination phase was used for analysis, yielding 1,200 temporal samples per trial at the 1,000 Hz sampling rate. For the visual task, a 1 s window starting at stimulus onset was extracted. Reports from several participants suggested reduced concentration toward the end of the 4 s trial, making the initial second the most reliable. This corresponded to 1,000 temporal samples per trial.

#### Data augmentation

To increase the size and variability of the MEG dataset while preserving class labels, we applied a simple temporal-shift augmentation to the source-reconstructed trials (auditory: 225 × 202 × 1,200; visual: 90 × 107 × 1,000; trials × source points × time points). For each trial we created two additional, time-shifted copies by shifting the signal forward by 100 ms and 200 ms, respectively. This procedure was inspired by the study of Dash et al. (2020). Shifting was implemented at the sample level (sampling rate 1,000 Hz) by removing the first *n* samples (corresponding to the desired delay) and padding zeros at the end of each trial. The original, 100 ms-shifted, and 200 ms-shifted trials were then concatenated, tripling the number of trials (auditory: 675; visual: 270). The class labels were duplicated accordingly so that each augmented trial retained the correct stimulus label. Such temporal jittering also accounts for intertrial variability in participants’ reaction times and imagery onset, thereby compensating for small temporal misalignments in the data while effectively increasing the dataset size. We thereby introduced small temporal variability, which can improve model robustness without distorting the underlying neurophysiological patterns.

#### Feature extraction

For each MEG trial (auditory and visual), we derived time–frequency representations using a continuous wavelet transform (CWT). We employed the Python library `scipy.signal.cwt` together with the Morlet wavelet `morlet2` to compute time–frequency representations of each MEG trial. The CWT convolves the signal with scaled and shifted Morlet wavelets, yielding complex coefficients whose magnitude reflects signal power across frequencies and time.

Single-channel signals were decomposed with complex Morlet wavelets spanning 1–120 Hz on a logarithmically-spaced grid of 200 center frequencies (sampling rate 1,000 Hz). The wavelet parameter was set to *w* = 6, balancing temporal and spectral resolution. The absolute values of the complex coefficients were retained to yield amplitude scalograms of shape (frequencies × time).

This procedure was applied independently to each source point signal, resulting in a set of channel-wise two-dimensional scalograms per trial. Inspired by the approach of Dash et al. [2020], the individual channel scalograms were spatially arranged into a single composite image representing the time–frequency activity across all sources in the ROI. The amplitude matrices were visualized using the matplotlib viridis colormap which maps low amplitude values to dark violet/blue and high amplitude values to yellow/green tones. These composite images were saved as 1000 × 1000 × 3 arrays, where the three channels corresponded to the RGB-encoded red, green, and blue components of the color-mapped amplitude values. This RGB encoding increased the dimensionality and complexity of the input, allowing the deep neural network to exploit richer feature hierarchies during training. These images were then further processed and further used as input to the employed models.

#### Model variants

Data were analyzed using a within-subject design, with separate models trained for each participant. Both ROIs described above were used to examine task-specific neural activity, resulting in two model groups. The first group used source-reconstructed data from the task-relevant cortical region (e.g., visual cortex for the visual task), ensuring that model inputs matched the functional demands of the condition. The second group used data from the non-task-relevant cortex (e.g., visual cortex for the auditory task). This setup enabled us to assess the specificity of neural representations and to reveal potential cross-modal activity.

#### Classification frameworks

To evaluate both simple and complex decision boundaries, we trained two complementary classifiers on the wavelet–scalogram features: a regularized linear model and a deep convolutional neural network. For the linear model, the scalogram images (1000 × 1000 × 3) were converted to single-channel grayscale, as the linear classifier requires one-dimensional feature vectors rather than multi-channel image data. The grayscale images were then resized to 256 × 256 pixels to reduce computational load, and z-scored per image. This simple linear architecture served as a baseline, assessing whether class information could be separated by a linear decision boundary in feature space. The model will, in the following, be referred to as LogReg. We performed a 70/15/15 split of data for training, validation and testing, respectively. To prevent data leakage, every original trial and its augmented copies (+100 ms and +200 ms time shifts) were assigned to the same set.

To capture more complex, non-linear relationships in the time–frequency representations, we used AlexNet [Krizhevsky et al., 2012]. AlexNet comprises stacked convolutional layers with non-linear activations and pooling, followed by fully connected layers. This hierarchical architecture learns progressively more abstract spatial features, from simple edges and frequency contours to complex joint time–frequency motifs, directly from the input images. Due to the MEG scalograms being effectively two-dimensional time–frequency images, AlexNet is particularly well suited to extract multiscale spatial structure and detect discriminative oscillatory patterns across sources and timescales. Importantly, Dash et al. [2020] previously demonstrated that AlexNet yields strong performance for imagined speech decoding, providing additional justification for its use in the present study. The model will, in the following, be referred to as CNN.

For input to the CNN, the scalograms were, as for the LogReg model, resized to 256 × 256 pixels and normalized. The network was initialized with ImageNet weights and fine-tuned employing the Adam optimizer using a learning rate of 0.001 and a batch size of 64. We applied the same grouping strategy to the train, validation, and test splits as for the linear classifier, guaranteeing that all augmented versions of a given trial remained within a single split. Training proceeded for 30 epochs.

All networks were implemented using the PyTorch framework in Python 3.12.2. Models were trained on an NVIDIA RTX A5000 GPU with 24 GB RAM and CUDA 12.6.

#### Statistical evaluation

To determine whether model accuracies were above chance and to directly compare models, we performed one-sample and paired-sample t-tests. For each participant and condition, the mean decoding accuracy was tested against the theoretical chance level of 33% (three-class problem) using a one-sample t-test. To assess whether one classifier outperformed another, we used a paired-sample t-test comparing their accuracies across participants. All tests were performed in the Python library scipy [Virtanen et al., 2020] and significance was assessed at *p* < 0.05.

The subject identity was added as random effect to account for within-subject correlations. Model coefficients, and *p*-values were extracted from the fitted models. By considering all combinations of classifier type (CNN or logistic regression), task (auditory and visual) and ROI (task-relevant or non–task-relevant), we obtained eight models in total. Therefore, *p*-values of the linear mixed-effect models were corrected for eight comparisons using Benjamini-Hochberg FDR correction.

## Results

### General model performances and regional specificity

The mean accuracies of the model for the auditory and visual decoding tasks are visualized in Figure 2. The LogReg models, as well as the CNN approach, were trained and evaluated once per participant. We trained one network per subject using functionally relevant data and a second model using data from the non-task-relevant ROI. The two investigated ROIs are shown in Figure 2a. This resulted in task-relevant models, namely for the auditory task on auditory cortex (for both LogReg and CNN) and for the visual task on visual cortex (for both LogReg and CNN). We also computed non-task-relevant models, namely for the auditory task on visual cortex (for both LogReg and CNN), as well as for the visual task on auditory cortex (for both LogReg and CNN).

**Figure 2.**
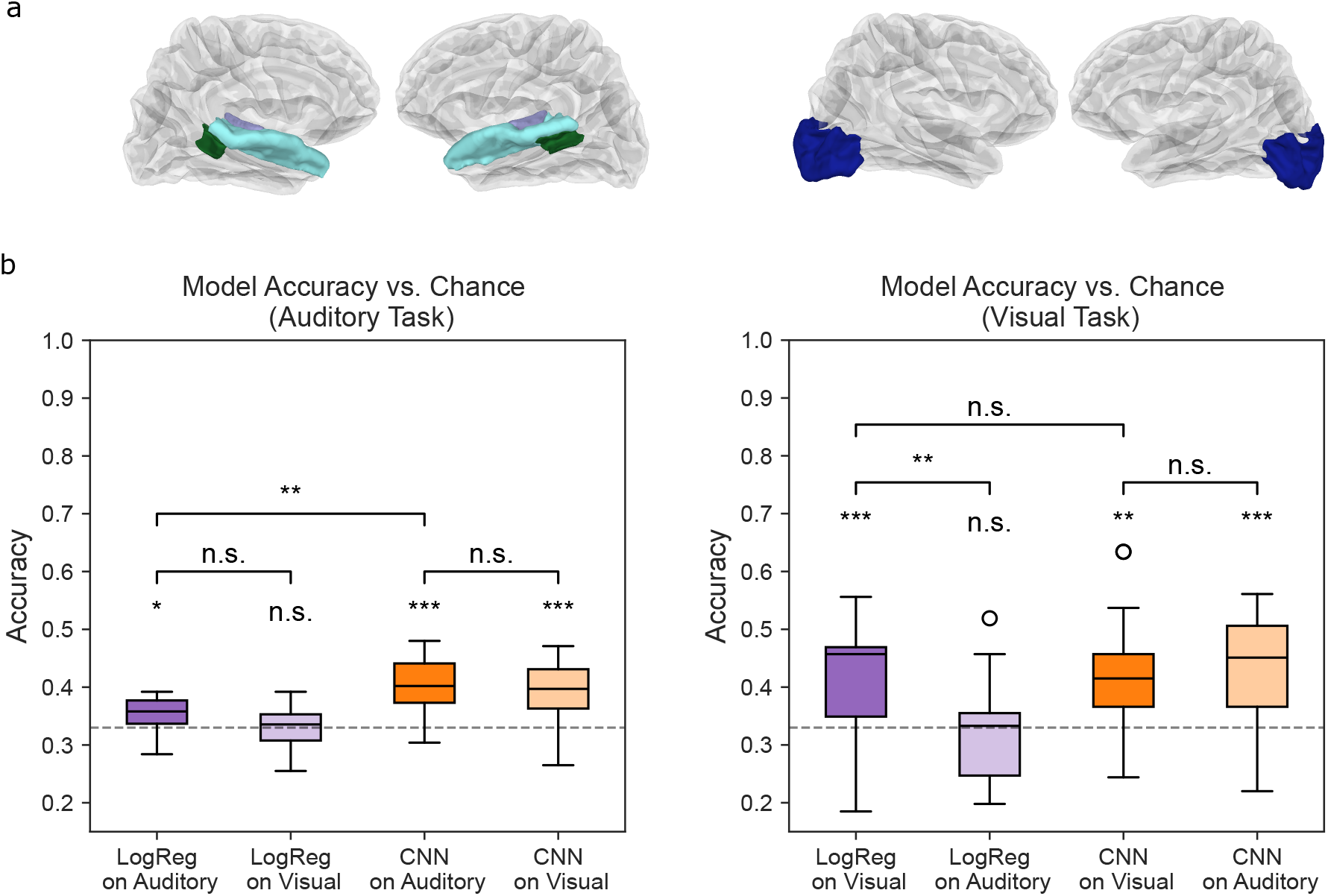
(a) Auditory and visual ROIs consisting of the transverse temporal gyri (purple), the superior temporal gyri (turquoise), the banks of the superior temporal sulci (dark green) in the auditory ROI (left) and the occipital lobe (dark blue) in the visual ROI (right). (b) Classification accuracies of the logistic regression model (LogReg, purple) and AlexNet (CNN, orange) models for auditory (left) and visual (right) decoding tasks. Dark colors indicate models trained on the task-relevant ROI (auditory cortex for the auditory task; visual cortex for the visual task), and light colors indicate models trained on the non-task-relevant ROI. Dashed lines mark the three-class chance level (33%). Significance stars denote one-sided t-tests against chance, as well as two-sided paired t-tests to compare model performances against each other (∗, *p* < 0.05; ∗∗, *p* < 0.01; ∗ ∗ ∗, *p* < 0.001, n.s. not significant).

Figure 2b demonstrates the classification accuracies of the CNN models and the LogReg models for the different conditions. Significance annotations indicate whether performance significantly exceeds chance. In Figure 2b, the classification accuracies of the CNN models and the LogReg models are compared to the chance level (33%, three-class classification). The left panel depicts the data from the auditory task, and on the right, the data from the visual task is presented. The performances of the LogReg models are shown in purple, the CNN accuracies in orange. The darker purple and orange depict accuracies of the models that were trained on the task-relevant ROI and the lighter purple and orange are the accuracies of the models which were trained on the respective non-task-relevant ROI. All models, including CNN instances and LogReg models, trained on data from the functionally relevant ROI (dark purple and dark orange), on average, significantly exceeded the chance level (33%). This was the case for both the auditory and the visual task (Figure 2b left and right panels). For the CNN models, this difference was found to be statistically significant with *p*-values less than 0.001 (auditory task) and less than 0.01 (visual task), respectively. The LogReg models also outperformed chance level with a *p*-value below 0.05 (auditory task) and less than 0.001 (visual task). Regarding the non-task-relevant ROIs, the LogReg model accuracies (light purple) did not exceed chance-level accuracy, whereas the CNN model accuracies significantly outperformed chance-level, with *p*-values less than 0.001 for both tasks.

Comparing the models trained on the task-relevant ROI with those trained on the non-task-relevant ROI, we only found a significant difference for the LogReg model of the visual task. Here, the model trained on the task-relevant ROI significantly outperformed the model trained on the non-task-relevant ROI (Figure 2b, *p* < 0.01).

### LogReg vs. CNN: performance evaluation

To compare the CNN-based approach with the linear approach, we contrasted model accuracies of LogReg against the CNN accuracies. We here concentrated on the models which were trained on the task-relevant ROI. As shown in Figure 2b, in the auditory task, performance was superior with the CNN compared to the LogReg model (*p* < 0.01). Conversely, in the visual task, the LogReg model did not outperform the CNN model.

### Model performances during auditory and visual tasks

When comparing the overall performance between data from the auditory versus visual imagery task, we observed that the models achieved either comparable or superior decoding for the visual than for the auditory imagery (Figure 3). While all models significantly outperform chance level, as it was already depicted in Figure 2b, the LogReg model for the visual task significantly outperformed the LogReg model for the auditory task. No such difference was found for CNN models when comparing auditory and visual tasks.

**Figure 3.**
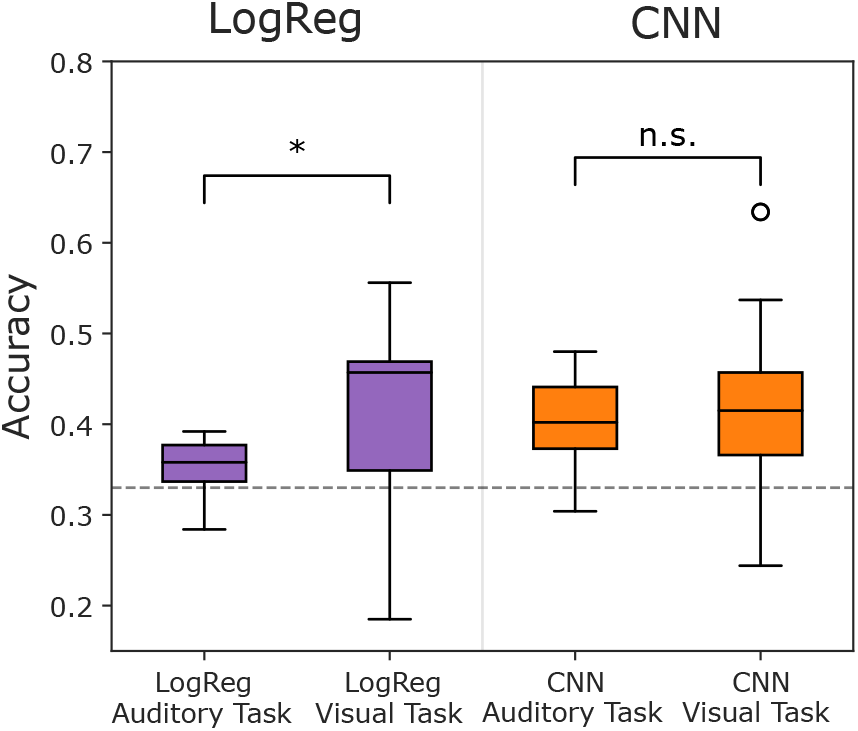
Comparison of decoding performances between auditory and visual imagination tasks. Mean classification accuracies of the logistic-regression models (LogReg, purple) and convolutional neural network (CNN, orange) models trained on task-relevant ROIs are shown. All models significantly exceeded the three-class chance level of 33%. The Lo-gReg model achieved significantly higher accuracy in the visual task than in the auditory task (∗ *p* < 0.05,), whereas no significant task-related difference was observed for the CNN models.

### Correlation between memorization task performance, VARK scores and model accuracies

After evaluating the performance of the models in general, we investigated correlations between the decoding accuracies and the additional behavioral data, which indicated memory capacity and individual learning preferences collected from the subjects. We therefore computed linear mixed-effect regression models, which show the dependency of the memorization task and *VARK* scores (*MT*_*a*_, *MT*_*v*_ and *VARK*_*a*_, *VARK*_*v*_, *VARK*_*k*_, *VARK*_*r*_) on the model accuracies. We did this analysis for the decoding accuracies of the LogReg as well as the CNN model for both tasks, auditory and visual, on the task-relevant ROI and also on the non-task-relevant ROI. Figure 4a depicts the results for the auditory and visual classification tasks with the data that was source-reconstructed in the task-relevant ROI (auditory task data on auditory cortex and visual task data two-sided paired t-test on visual cortex). The memorization task performances *MT*_*a*_ and *MT*_*v*_ and the *VARK* scores were tested for dependency on the LogReg accuracies (upper panel) and on the CNN accuracies (lower panel). For the auditory task, we found a significant (*p* < 0.05) positive dependency between LogReg accuracies and the auditory *VARK*_*a*_ score. Moreover, the CNN accuracies for the auditory task showed a significant (*p* < 0.001) negative link with the visual memorization task scores and a significant (*p* < 0.05) positive link with the kinesthetic *VARK*_*k*_ score. Regarding the visual task in the task-relevant ROI (Figure 4 a, lower left panel), a tendency for a positive association between the visual memorization task score (*MT*_*v*_) and the CNN decoding accuracies can be observed (*p* < 0.05 before correction for multiple comparisons; the *p* value is above 0.05 after correction). The respective results for the auditory and visual classification tasks with the data reconstructed in the non-task-relevant ROI (auditory task data on visual cortex and visual task data on auditory cortex) can be seen in Figure 4b. Here, no significant influence of behavioral data on the model performance can be observed, neither regarding the auditory task nor the visual task.

**Figure 4.**
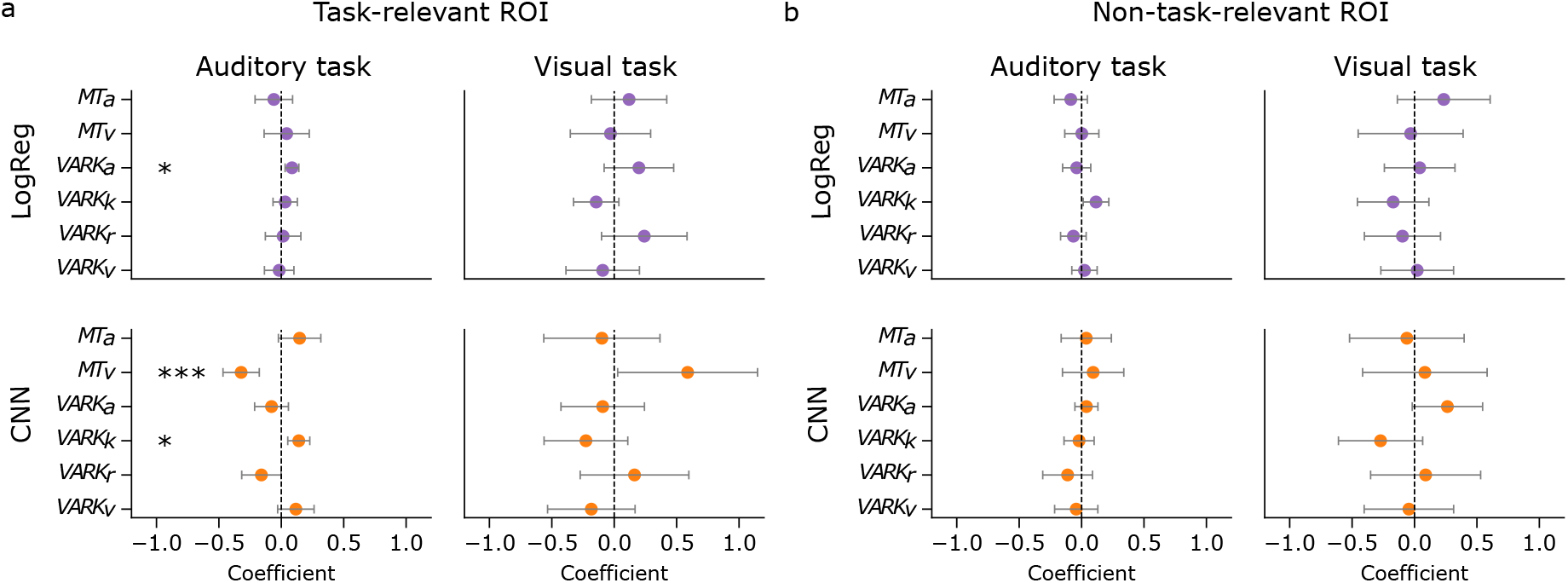
Relationship between behavioral measures and decoding accuracies. (a) Results of linear mixed-effects regression models for the auditory and visual classification tasks in the respective task-relevant ROI: model coefficients (± standard deviation of the mean) for the influence of auditory and visual memorization task performance (*MT*_*a*_, *MT*_*v*_) and *VARK* subscores (*VARK*_*a*_, *VARK*_*v*_, *VARK*_*k*_, *VARK*_*r*_) on the logistic regression (LogReg, upper panel) and CNN (lower panel) accuracies for the auditory task (left) and the visual task (right). Significance is indicated by markers: ∗ *p* < 0.05 and ∗ ∗ ∗ *p* < 0.001, all corrected for 8 comparisons. (b) Results for the auditory and visual classification tasks in the resepective non-task-relevant ROI.

## Discussion

### General model performances and regional specificity

Linear logistic regression models trained on task-relevant cortical regions, i.e., auditory cortex for auditory imagery and visual cortex for visual imagery, consistently achieved higher accuracies than their respective control models trained on non-task-relevant ROIs. This confirms that modality-specific regions provide the most informative neural features for imagery decoding. However, a striking observation in this study is that the CNN (AlexNet) achieved a decoding performance that was much above chance, for both the auditory and visual tasks, even when trained on non-task-relevant regions. For the visual task, the CNN trained on the auditory cortex even tended to outperform the model trained on the visual cortex. This indicates that the neural representations underlying imagery are not strictly localized, but instead distributed across partially overlapping cortical networks. Nishimura et al. (2015) reported that visual thinkers activate visual cortical regions even during verbal tasks, indicating that visual processing can accompany language-related activity. Such overlapping and individually variable activation patterns may explain why decoding succeeded even in non-task-relevant regions and why the CNN models, in particular, were sensitive to these distributed neural representations. Additionally, several experimental factors may contribute to this finding. During the auditory task, participants viewed the written cue on the screen and were instructed to keep their eyes open and fixate on a central cross throughout the imagery period. This sustained visual engagement likely activated occipital regions, which may have carried information correlated with the imagery process and thus enabled decoding from the visual cortex. In contrast, during the visual task, participants might have silently labeled the presented images (“Herz,” “Stern,” “Baum”), introducing activity in auditory or language-related areas.

Together, these effects could explain why decoding performance was above chance even in non-task-relevant regions and why cross-modal contributions emerged in the CNN models. Unlike the linear classifier, which relies on a single global decision boundary, the CNN can capture non-linear and spatially distributed feature patterns that may jointly encode information from multiple sensory systems. In this context, subtle and spatially overlapping activations in visual and auditory cortices, arising from visual fixation during the auditory task or verbal labeling during the visual task, are likely too weak or heterogeneous to be linearly separable. However, the hierarchical convolutional layers of the CNN can integrate these small, distributed signals across sources and time–frequency scales, effectively exploiting multimodal information that the linear model cannot. The results are consistent with previous evidence for functional overlap between visual and auditory processing systems, where visual areas are recruited during inner speech and verbal thinking [Amit et al., 2017]. The ability of deep neural networks to exploit such distributed, cross-modal activity patterns underscores their potential for decoding complex cognitive states that engage multiple sensory systems simultaneously.

### LogReg vs. CNN: performance evaluation

The comparison of a simple linear classifier (LogReg) with a CNN-based approach (AlexNet) revealed modality-dependent advantages. In the auditory task, the CNN consistently outperformed LogReg, reflecting its ability to capture complex, high-dimensional time–frequency features. This result aligns with earlier work demonstrating the suitability of AlexNet for imagined speech decoding [Dash et al., 2020]. In contrast, for visual imagery, the linear model tended to perform better than the CNN in the task-relevant visual ROI. One explanation is that the auditory dataset was higher-dimensional with more source points (202 in the auditory ROI, 107 in the visual ROI) and longer time windows (1.2 s for the auditory task and 1 s for the visual task), making it more suitable for hierarchical feature learning, whereas the smaller visual dataset was more easily linearly separable. Moreover, the auditory task comprised the imagined articulation of a spoken word, which is limited in duration, leading to unavoidable noise before and after the imagined phase. In contrast, the visual task was the imagination of a picture, which easily filled the whole 4 s imagery phase window, leading to potentially higher signal-to-noise ratio and more stable neural representations. Higher-dimensional data can introduce greater complexity, which may benefit deep learning models like AlexNet that leverage hierarchical feature extraction [LeCun et al., 2015]. In contrast, lower-dimensional datasets often favor linear classifiers such as logistic regression models, as they are more likely to be linearly separable [Hastie et al., 2009]. Additionally, high-dimensional feature spaces can hinder the performance of linear models due to the curse of dimensionality [Bishop and Nasrabadi, 2006], making it more challenging to effectively separate classes. On the other hand, lower-dimensional datasets pose a higher risk of overfitting for deep neural networks, as they may not contain sufficient variability to generalize well to unseen data [Hastie et al., 2009]. These modality-specific differences in data structure and task design likely explain why the CNN benefited more from the auditory dataset, whereas the linear model was sufficient to exploit the more robust representations present in the visual condition. Together, these findings emphasize that the choice of classifier should be guided by both the dimensionality of the data and the temporal characteristics of the imagery task.

### Model performance across tasks

Overall, decoding accuracies were higher in the visual task compared to the auditory condition, particularly and significantly for the LogReg model. Several factors likely contribute to this difference. Visual imagery benefits from short-lived iconic memory traces that maintain perceptual information after stimulus offset [Sperling, 1960], whereas auditory imagery requires active reconstruction of the stimulus, making it more demanding. Kilmarx et al. (2024) found that visual imagery following direct visual presentation produced stronger EEG signatures than spontaneous imagery from memory, suggesting that short-term, cue-based visual imagery elicits robust and classifiable neural responses. The sustained visual engagement during our trials may have similarly facilitated stable neural representations and enhanced model performance.

Moreover, visual imagination occurred within the same modality as stimulus presentation (image-to-image), whereas auditory imagination required a cross-modal transformation from visual cues to internal auditory representations. Interestingly, CNN performance did not differ significantly between tasks, suggesting that deep models are less sensitive to such modality-specific constraints. This may reflect their ability to extract subtle, distributed features that transcend modality boundaries, thereby equalizing performance across auditory and visual conditions. The observation that CNNs could decode imagery from non-relevant regions further supports the idea that imagined perception involves widespread cortical engagement, possibly mediated by top-down feedback between sensory areas. Such cross-modal representational overlap may also imply that combining auditory and visual imagery in future paradigms could enhance decoding performance by exploiting these shared neural substrates.

### Correlation between memorization task performance, VARK scores, and model accuracies

Beyond general decoding performance, exploratory analyses revealed meaningful associations between behavioral measures and model accuracies. For the auditory task, higher LogReg accuracies were associated with stronger auditory learning preferences (*VARK*_*a*_), indicating that participants who naturally process information auditorily produced more discriminable neural representations of imagined speech. In contrast, the CNN revealed a different pattern: its accuracies for the auditory task were positively correlated with kinesthetic learning preference (*VARK*_*k*_) and negatively correlated with visual memory performance (*MT*_*v*_). These effects suggest that participants who rely less on visual strategies, or who engage more embodied or motor-related representations, may generate richer or more distributed auditory neural patterns that deep models can exploit. On the other hand, the statistically insignificant association but strong positive tendency between the CNN performance of the visual task and the visual memorization task score indicates that participants with stronger visual memory capacities appear to form clearer and more stable neural patterns during visual imagination, whereas participants with weaker visual memory capacity tend to perform better in the auditory task. Studies indicate that individuals who classify themselves as visual learners tend to have enhanced understanding and recall of newly acquired information, which contributes to better outcomes in evaluative tasks [Jawed et al., 2024]. The fact that behavioral correlations emerged for only one model (LogReg or CNN) at a time suggests that each classifier type may capture partly different aspects of the underlying neural signal. The linear model appears to reflect more direct, modality-specific associations in the auditory task, such as auditory preference improving imagined speech decoding, whereas the CNN, by modeling more complex signal patterns, may emphasize other dimensions of individual variability. Importantly, the absence of overlapping effects between models indicates that these relationships are not driven by subject-individual learning preferences or task performances, but by how each architecture extracts information from the MEG data. This model-specific sensitivity underlines that behavioral–neural correlations should be interpreted carefully in light of the decoding approach used, as different algorithms may highlight distinct facets of the same cognitive process. Notably, no significant behavioral effects were found in models trained on non-task-relevant regions, suggesting that while cross-modal representations enable above-chance decoding, the behavioral relevance of these signals is much stronger within the task-relevant cortical regions.

### Limitations and conclusion

Although the classification accuracies were above chance level, it must be acknowledged that the overall decoding performance remained relatively low, which could be due to several different limitations in our work. One potential aspect impeding classification performance, particularly during the auditory task, could be the challenge of accurately identifying the timing of the imagined event. Temporal misalignment in the recorded neural signals can introduce noise, potentially reducing the effectiveness of the classification models. We partly accounted for this by cutting the data in the auditory task from 2.5 s duration to only 1.2 s duration and in the visual task from 4 s to 1 s, selecting the time intervals when mental imagery was assumed to be most pronounced. However, since the length and, thus, articulation duration of the words (*Absicht, Idee*, and *Ehre*) differ from subject to subject and even between trials, especially the data where the shorter words had to be imagined, are likely to still contain significant noise.

A further limitation concerns participant fatigue and attentional fluctuations. The one-hour task duration was demanding, and no direct behavioral measures of concentration were collected in the breaks. Incorporating brief random control checks or task-related recall prompts, as suggested by Nieto et al. (2022), could help track engagement and improve data quality in future studies.

One should keep in mind that the number of participants in the study (*n* = 18) may further limit the power to detect statistical significance, making it more challenging to identify meaningful correlations or effects, especially when it comes to individual learning preferences. A larger sample size in a future study may provide more statistical power, potentially revealing a correlation in the auditory task. A larger number of subjects would also increase the available amount of training data for the decoding algorithm, presumably further boosting mental imagery detection.

Our findings advance the understanding of how cognitive processes such as memory and imagination are represented in the brain and demonstrate the potential of modern deep learning for their decoding. The CNN (AlexNet) achieved robust above-chance performance across modalities and regions, suggesting that deep models can exploit distributed neural information and are less dependent on precise anatomical targeting. This robustness may prove valuable for future BCI development, where recording coverage and user-specific variability often limit decoding performance. Finally, the observed relationship between individual learning preferences and decoding performance points toward the possibility of personalized BCIs that adapt to a user’s cognitive strengths, emphasizing visual or auditory channels according to individual processing preferences. Such approaches could enhance the efficiency, robustness, and accessibility of next-generation BCI systems.

## Data Accessibility

The data used in this study is available from the authors upon request.

## Acknowledgments

This project was supported by the German Federal Ministry of Education and Research (Cluster4Future SE-MECO, project number 03ZU1210FB) and the German Science Foundation (DFG, project number 523344822).

## Author contributions

AS and TR designed research; AS performed research; AS and JR collected the data; AS, MS, CJ and TR analyzed data and wrote the paper.

## Conflict of Interest

The authors declare no conflict of interest.

## Notes

### Competing Interest Statement

The authors have declared no competing interest.

